# Undergraduate GPA does not predict success in PhD programs for cohorts of MS students at two minority-serving institutions

**DOI:** 10.1101/2025.01.03.630681

**Authors:** M Fuse, K Rath, B Riggs, RM Esquerra, A Peterfreund, C Gutierrez, F Bayliss

## Abstract

Master of Science (MS) research training programs funded by organizations such as the NIH, the NSF as well as private corporations represent the potential for significant interventions for student success, especially for marginalized or underrepresented and first-generation students, to bridge the gap between undergraduate studies and doctoral programs. These students often face serious challenges during their undergraduate years, such as navigating unfamiliar academic systems, balancing demanding coursework with work responsibilities, and fulfilling family obligations. These and other systemic pressures can impede their academic progress and opportunities for research experience. We therefore asked how their undergraduate GPA impacted success in the PhD in terms of (i) acceptance, (ii) completion, and (iii) time to degree after participating in a funded MS research training program. We examined data collected at San Francisco State University and California State University, Los Angeles over a 30-year period because they (i) had similar student demographics, (ii) were institutions with strong MS degrees, and (iii) had an infrastructure (established offices) to coordinate various training programs. We found that high GPA did not predict greater success in entering or completing the PhD, or in time to degree. This study, therefore, demonstrates that undergraduates from diverse groups with lower undergraduate GPA levels can benefit from a structured MS research training program in reaching their goal of becoming a PhD scientist. Moreover, it indicates that more holistic approaches in admissions are required, including limiting the use of a GPA below 3.0 as an early filter to eliminate applicants. These MS programs appear to play a crucial role as a bridge to PhD studies and are instrumental in enhancing diversity within STEM fields. We suggest that a strategy for success is to provide infrastructure in the form of a coordinated office housing the training grants as a means of structuring mentorship and professional development for under-represented and first generation students.

## 1. Introduction

Graduate admission committees seek to identify curious, creative, and motivated candidates with the academic preparation necessary to complete the PhD and pursue productive research careers in academia, scientific industry, and other sectors of the science enterprise. Committees depend on a review of application materials that generally include letters of recommendation, statements of purpose, descriptions of prior research experiences, standardized test scores, and transcripts of undergraduate coursework. Historically, undergraduate grade point average (GPA) has been a significant criterion, presumably as a measure of academic preparation for graduate study. Undergraduate GPA, while usually considered a measure of academic performance, can sometimes reflect a student’s level of privilege and access to resources, including the ability to focus exclusively on studies without balancing work or other responsibilities. Here, we argue that undergraduate GPA can be a limited and often misleading indicator of a student’s talent, motivation, and readiness for graduate study. As such, it may serve as an unnecessary and potentially exclusionary filter.

Students from historically marginalized communities often face significant barriers in pursuing PhDs, which include limited access to educational opportunities, financial constraints, lack of representation and mentorship in academia, systemic biases, and discrimination within higher education institutions. Discrimination and biases in gatekeeping practices for entry into PhD programs include selection criteria based on undergraduate GPA or graduate record exam (GRE) scores, where low scores are assumed to indicate an inability to succeed. Over the past decade, many discussions have involved which criteria in PhD applications can predict a student’s success in doctoral programs. It is becoming increasingly clear that quantitative metrics such as GRE and GPA scores fail to predict success (Miller and Stassun, 2014; Wolf, 2014; Weiner, 2014; Hall *et al*., 2017; Moneta-Koehler *et al*., 2017; Petersen *et al*., 2018; Miller *et al*., 2019; Sealy *et al*., 2019; Wilson *et al*., 2019; Mendoza-Sanchez *et al*., 2022). They do not appear to predict success in the outcomes most valued by the PhD community: completing the PhD program, research productivity, and career success in research. More problematic are charges that these metrics exhibit cultural and gender biases in outcomes, and select against socioeconomically disadvantaged populations (Miller and Stassun, 2014; Wolf, 2014; Posselt, 2016; Hall *et al*., 2017; Moneta-Koehler *et al*., 2017; Petersen *et al*., 2018; Wilson *et al*., 2018; Wilson *et al*., 2019).

An additional concern with GPA is grade inflation, where undergraduate GPAs have risen over the decades without a corresponding increase in actual academic achievement. Although grade inflation has been reported across all sectors of higher education, it has been observed at higher rates among highly selective private colleges over the past 30 years and at lower rates in less selective commuter public institutions (Rojstaczer, 2010; Rojstaczer, 2012). Even within the same institution, trends show higher grade inflation among students from higher socioeconomic backgrounds. At one exclusive college, students from families earning $250,000 or more were 60% more likely to have significantly higher GPAs than their counterparts from families with incomes of $40,000 or less (Bikales, 2022).

Recently, a more holistic approach has involved evaluating more characteristics, including research experiences, letters of recommendation, and the statement of purpose. For instance, a study at the University of North Carolina found that higher ratings from recommendation letters correlated with publication output (Hall *et al*., 2017). Likewise, the years of prior research experience correlated with the highest and lowest-ranked students within the University of California San Francisco (UCSF) graduate student population (Wiener, 2014). Because of poor correlation between these metrics and desired graduate program and career outcomes, many major research universities have removed general GRE scores as requirements for PhD applications, and a more holistic application approach has been taken as a strategy for mitigating the bias involved in the review of applicants (Langin, 2022). However, undergraduate GPA continues to be used as an initial cutoff for applications, typically set at 3.0. In this regard, it should be noted that, in many of the studies cited above, analyses were conducted on PhD cohorts that had already been pre-selected based on an undergraduate GPA cutoff of 3.0.

We had the opportunity to examine cohorts of under-represented students in PhD programs over a 30-year period, whose undergraduate GPA levels ranged from 2.0 to 4.0 and who entered research-focused Master’s degree programs before entry to PhD studies. Our goal was to determine whether undergraduate GPA could predict success in PhD programs for students who participated in a funded research-based Master’s training program designed to provide research experience and academic preparation for doctoral studies. We looked at PhD outcomes from students who attended two masters serving institutions from the California State University (CSU) system and participated in (primarily) NIH-funded research training programs. Student outcomes from these CSU programs, at San Francisco State University (SFSU) and California State University, Los Angeles (CSULA), were examined for student admission and completion of doctoral training programs. As one of the largest public university systems in the United States, CSU serves a student body that is the most diverse in the nation in terms of ethnicity, economic background, and age (California State University Introduction, 2023; SFSU Institutional Research), allowing for a large sample size of under-represented students in these programs. At SFSU, the Student Enrichment Opportunities (SEO) office oversaw students funded by federal and private training programs. At CSULA, the equivalent occurred in the Minority Opportunities in Research (MORE) office. In the past 30 years, over 95% of students participating in these diversity training programs at SFSU and CSULA were underrepresented (UR), as defined by the National Institutes of Health (NIH), which includes Black or African American, Hispanic or Latino, American Indians or Alaskan Natives, Native Hawaiians and other Pacific Islander groups, individuals with disabilities, and individuals from disadvantaged backgrounds. Most also met other criteria for underrepresentation, including being low income and first-generation university students.

For this study, we examined the records of three decades (1992-2019) of students in Master of Science programs at SFSU and CSULA who participated in research training programs designed to prepare them for admission to and success in strong PhD programs. These research training MS-to-PhD programs were supported by grants from the NIH, NSF or private foundations. After the MS, over 85% of the students who applied to PhD programs successfully gained admission to and completed strong PhD programs across the country.

In our MS-to-PhD training programs, we collected all the documents required of students applying to PhD programs: letters of recommendation, statement of purpose, description of prior research experience, the general GRE, and transcripts of undergraduate coursework. When PhD programs discontinued using the general GRE, we also no longer required these scores. Student selection for these programs was strongly dependent on a holistic review of these documents along with extensive in-person interviews for each candidate. These conversations revealed much about the applicants’ curiosity, rigor of undergraduate coursework, motivation for research careers, and aptitude for this venture. Typically, there was a significant number of applicants who demonstrated clear strengths in curiosity about their scientific disciplines and interest and experience in research, yet whose undergraduate academic GPA was less than 3.0. These candidates were frequently among those highly recommended to us by faculty colleagues at our home and other institutions as talented individuals who had worked as volunteers in their research groups, an incongruity between these applicants’ undergraduate GPAs and their perceived talents and motivation.

Nevertheless, all candidates (including those with high GPAs) were required to demonstrate competence in the undergraduate content areas of their major scientific disciplines. Some departments administer to all incoming MS students diagnostic placement examinations developed locally, or by their national scientific societies. At times, a student that demonstrated deficiency in a necessary undergraduate science content area was given opportunity to address this prior to registering in an MS course requiring that content knowledge through self-study and retesting in the appropriate content area, or auditing the appropriate undergraduate course. However, it is noteworthy that all students admitted with undergraduate GPAs less than 3.0 maintained a solid level of academic achievement throughout their MS programs indistinguishable from that of students admitted with higher GPAs (data not shown).

Given the typical undergraduate GPA cut-off of 3.0 for many R1 institutions, we wanted to see if there were differences in success for participating Master’s students with low undergraduate GPAs (<3.0) versus high undergraduate GPAs (≥3.0), as measured by (i) the number of students matriculating as PhD students, (ii) the completion rates, and (iii) the time to degree in the PhD programs. This was particularly significant, given that typically, only half of the students who enter STEM doctoral programs in the US will graduate with a doctoral degree (NCSES NSF, 2022). Our study found that a high undergraduate GPA did not predict success in entering or completing a PhD program, nor did it predict time to degree. These findings suggest that a more holistic review of applicants - one not centered on undergraduate GPA and that can include the quality of the training program the students experienced - is a better mechanism for predicting student success in doctoral programs. We also provide an example of how such MS training grant programs enhance their institutions and discuss the potential role they and their affiliated offices may play in facilitating student success.

## 2. Methods

To assess three decades of student entrance into PhD programs (1992-2019 for SFSU; 1998-2018 for CSULA), we took advantage of the coordination of training programs in the offices housing these grants at SFSU-SEO and CSULA-MORE. There was a concerted effort by the two offices to track all students through multiple sources, including LinkedIn, emails, phone calls, and PhD institution web pages. At SFSU, we identified the educational outcomes for 330 master’s students supported by SEO programs who applied to PhD programs, between the fall of 1992 and the fall of 2019. Of the 330 scholars, 293 entered a PhD program (88.8%). At CSULA, as part of the MORE programs, 176 students were identified between the fall of 1998 and the spring of 2018. Of the 176, 154 entered PhD programs (87.5%). We considered several variables important for our analysis, which included: (1) the undergraduate institution the student came from (more specifically, whether a student came from their own MS institution as an undergraduate or from an external undergraduate institution) to eliminate institutional bias; (2) the undergraduate GPA (UG GPA); (3) whether or not the student entered a doctoral program (medical and other health-related degree programs were excluded); and (4) time to complete the doctoral program. We assessed differences in success with PhDs (acceptance, completion, and time to degree) for all SEO and MORE students but also considered whether there was a difference if students came from SFSU/CSULA or another institution as undergraduate students for their respective MS degrees (Fig. 1). Differences were assessed by Chi-squared analysis.

**Figure 1.**
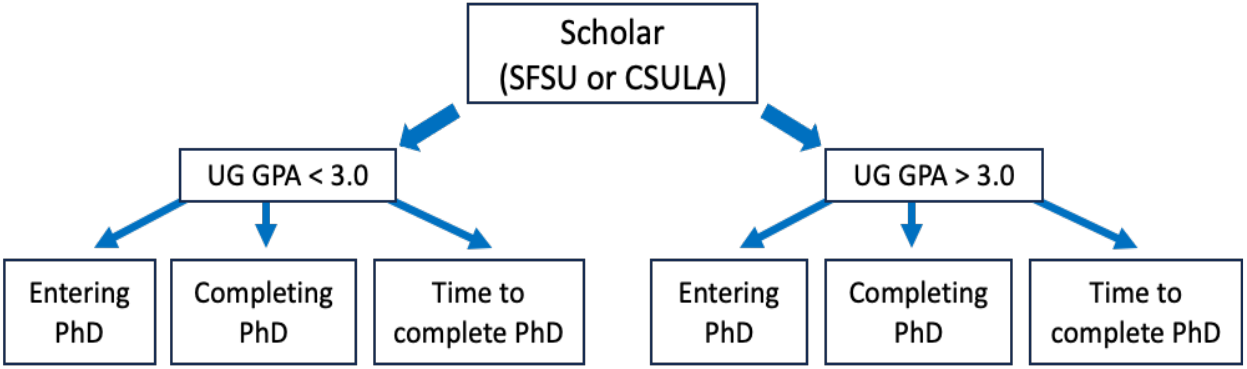
Flowchart of data collection from SFSU and CSULA.

Finally, we considered the change in the number and demographics of SFSU’s Biology MS students (including non-SEO students) over four decades as an example of the impact of training grants on the diversity of the student body within a STEM MS program. We collected student data from the SFSU Graduate Division to confirm the completion and graduation from the MS program for all Biology MS students from 1980 to 2019.

## 3. Results

### 3.1. Demographic characteristics of the MS student cohorts

Our data originated from over three decades of student participation in research training programs. We partitioned our cohorts based on undergraduate GPA (low UG GPA vs high UG GPA), whether they attended the MS institution as an undergraduate or another institution. For all of the analyses performed, we divided our MS students into two groups: Low UG GPA < 3.0 and High GPA ≥ 3.0 (Table 1). We considered whether they had completed their undergraduate degree at the same or another institution. Notably, our students were relatively evenly split between high and low UG GPA groups, with most scholars from Biology and Chemistry/Biochemistry. GPA data ranged from 2.0-4.0, with details in Supplemental Information Table S1.

**Table 1:** Demographics of MS students by university (SFSU or CSULA)

| Institution | SFSU | CSULA |
| --- | --- | --- |
| Total in sample | 330 | 176 |
| Low GPA (< 3.0) | 45.2% | 42.0% |
| High GPA (≥ 3.0) | 54.8% | 58.0% |
| Undergraduate at same institution | 43.0% | 33.5% |
| Undergraduate at another institution | 57.0% | 66.5% |
| Major: Biology | 79.4% | 42.6% |
| Major: Chemistry or Biochemistry | 14.2% | 31.3% |
| Major: Psychology | NA | 19.9% |
| Major: Other | 6.4% | 6.3% |

### 3.2. A high undergraduate GPA does not predict enhanced acceptance into a PhD program

In our first analysis, we looked to see if there was a difference between the two UG GPA groups in terms of whether they were accepted/entered a doctoral program or not for either SFSU or CSULA (Fig. 2; Tables 2 and 3, respectively). All groups showed high matriculation into PhD programs at 86-91% (Fig. 2). There was also no significant difference between low and high UG GPA groups (Tables 2 & 3). This was true when considering all students, or when separating them by the undergraduate institution they attended (P>0.05).

**Table 2:** Proportion of SFSU students from each UG GPA group entering a doctoral program, for all SFSU students and sorted by undergraduate university.

| Entering PhD | All SFSU |  | SFSU UG |  | non-SFSU UG |  |
| --- | --- | --- | --- | --- | --- | --- |
| % (n) | Low GPA (< 3.0) | High GPA (≥ 3.0) | Low GPA (< 3.0) | High GPA (≥ 3.0) | Low GPA (< 3.0) | High GPA (≥ 3.0) |
| Unknown | 0% (0) | 0.4% (1) | 0% (0) | 1.2% (1) | 0% (0) | 0% (0) |
| PhD program | 91.3% (136) | 86.7% (157) | 86.4% (51) | 83.1% (69) | 94.4% (85) | 89.8% (88) |
| Alternate | 8.7% (13) | 12.7% (23) | 13.6% (8) | 15.7% (13) | 5.6% (5) | 10.2% (10) |
| χ<sup>2</sup> test | 2.201 |  | 0.859 |  | 1.381 |  |
| p-value | 0.333 |  | 0.651 |  | 0.24 |  |

**Table 3:**
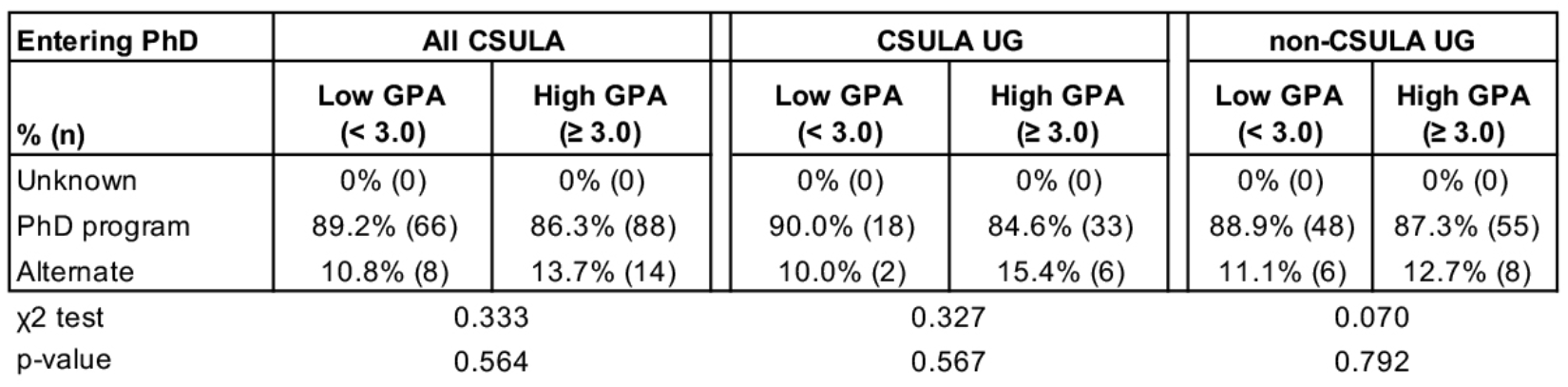
Proportion of CSULA students from each GPA group entering a doctoral program, for all CSULA students, and sorted by undergraduate university.

| Entering PhD | All CSULA |  | CSULA UG |  | non-CSULA UG |  |
| --- | --- | --- | --- | --- | --- | --- |
| % (n) | Low GPA (< 3.0) | High GPA (≥ 3.0) | Low GPA (< 3.0) | High GPA (≥ 3.0) | Low GPA (< 3.0) | High GPA (≥ 3.0) |
| Unknown | 0% (0) | 0% (0) | 0% (0) | 0% (0) | 0% (0) | 0% (0) |
| PhD program | 89.2% (66) | 86.3% (88) | 90.0% (18) | 84.6% (33) | 88.9% (48) | 87.3% (55) |
| Alternate | 10.8% (8) | 13.7% (14) | 10.0% (2) | 15.4% (6) | 11.1% (6) | 12.7% (8) |
| χ<sup>2</sup> test | 0.333 |  | 0.327 |  | 0.070 |  |
| p-value | 0.564 |  | 0.567 |  | 0.792 |  |

**Figure 2:**
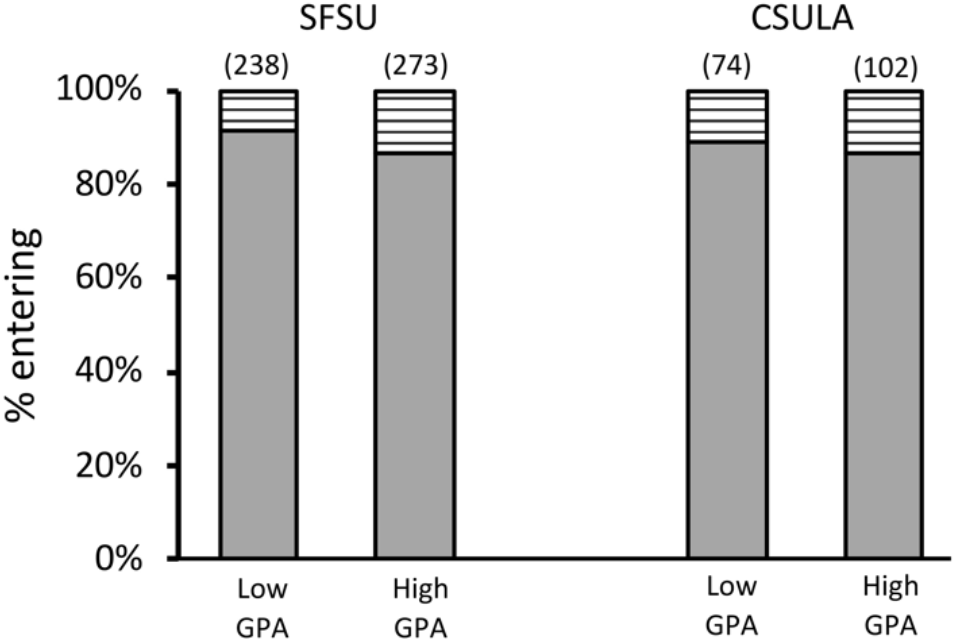
Proportion of SFSU or CSULA students from each UG GPA group entering a doctoral program (Solid bar). Total sample sizes are noted above the bars, in parentheses.

The range of PhD institutions accepting students into their programs was broad but similar for the low and high UG GPA groups (Tables S2, S3). Of the 293 SFSU students matriculating in PhD programs, the top five universities accepting SFSU students from the low UG GPA group were UC Davis, UC Berkeley, University of Washington, UCSF, and Harvard University. The top five universities accepting SFSU students from the high UG GPA group were UC Davis, UCSF, UC Berkeley, UCLA, and Harvard University (Table S2). Of the 154 CSULA students matriculating in PhD programs, the top five universities accepting students from the low UG GPA group were UCLA, UC San Diego, UC Riverside, UC Davis, UC Irvine/University of Illinois Urbana-Champaign (tied). The top five universities accepting students from the high UG GPA group were UCLA, University of Southern California, UC Riverside, UC Irvine, and. Claremont Graduate University (Table A3).

When we pooled all students so that UG GPA was not considered, there was also no significant difference for MS students getting into a PhD program based on where they completed their undergraduate degree (Table 4; P>0.07).

**Table 4:** Proportion of all students from their own or other institutions entering a doctoral program, sorted by undergraduate university and not by GPA.

| Entering PhD | UG Institution |  | UG Institution |  |
| --- | --- | --- | --- | --- |
|  | SFSU UG | non-SFSU UG | CSULA UG | non-CSULA UG |
| Unknown | 0.7% (1) | 0% (0) | 0% (0) | 0% (0) |
| PhD program | 84.5% (120) | 92.0% (173) | 86.4% (51) | 88.0% (103) |
| Alternate | 14.8% (21) | 8.0% (15) | 13.5% (8) | 12.0% (14) |
| $\chi^2$ test | 5.277 | | 0.091 | |
| p-value | 0.071 |  | 0.763 |  |

### 3.3. A high UG GPA does not predict enhanced completion of the PhD

We further assessed the PhD completion rate of the students in these two cohorts. At SFSU (Fig. 3), of the 293 MS students that entered PhD programs, there was no significant difference in terms of their ability to complete a PhD whether a student came into the SFSU research training program with a low or high UG GPA. This was true whether they completed their undergraduate degree at SFSU or elsewhere (Table 5). In contrast, a significantly greater proportion of the 154 CSULA MS students who completed their degrees came from the *low* UG GPA group compared to the high group (Fig. 3, asterisk). This significance was based on students completing their undergraduate degrees at a non-CSULA institution (Table 6, **bolded values**). That is, undergraduates from outside CSULA who entered the CSULA MS program were more likely to complete the PhD when they had a low UG GPA than a high one.

**Table 5:** Proportion of SFSU students from each GPA group completing, a doctoral program, sorted by undergraduate university. Those still in progress are noted below the p-values.

| Completing PhD | All SFSU |  | SFSU UG |  | non-SFSU UG |  |
| --- | --- | --- | --- | --- | --- | --- |
| % (n) | Low GPA (< 3.0) | High GPA ($\geq 3.0$) | Low GPA (< 3.0) | High GPA ($\geq 3.0$) | Low GPA (< 3.0) | High GPA ($\geq 3.0$) |
| Did not complete | 14.6% (15) | 14.5% (19) | 12.9% (4) | 21.2% (11) | 15.3% (11) | 10.1% (8) |
| Completed | 85.4% (88) | 85.5% (112) | 87.1% (27) | 78.8% (41) | 84.7% (61) | 89.9% (71) |
| $\chi^2$ test | 0.000 | | 0.893 | | 0.909 | |
| p-value | 0.990 |  | 0.345 |  | 0.340 |  |
| In progress | 33 | 26 | 20 | 17 | 13 | 11 |

**Table 6:** Proportion of CSULA students from each GPA group completing a doctoral program, sorted by undergraduate university. Those still in progress are noted below the p-values.

| Completing PhD | All CSULA |  | CSULA UG |  | non-CSULA UG |  |
| --- | --- | --- | --- | --- | --- | --- |
| % (n) | Low GPA (< 3.0) | High GPA ($\geq 3.0$) | Low GPA (< 3.0) | High GPA ($\geq 3.0$) | Low GPA (< 3.0) | High GPA ($\geq 3.0$) |
| Did not complete | 9.8% (6) | 21.1% (15) | 20.0% (3) | 14.3% (4) | 6.5% (3) | 28.9% (13) |
| Completed | 90.2% (55) | 78.9% (56) | 80.0% (12) | 85.7% (24) | 93.5% (43) | 71.1% (32) |
| $\chi^2$ test | 4.229 | | 0.234 | | 7.853 | |
| p-value | 0.040 |  | 0.629 |  | 0.005 |  |
| In progress | 5 | 17 | 3 | 5 | 2 | 10 |

**Figure 3:**
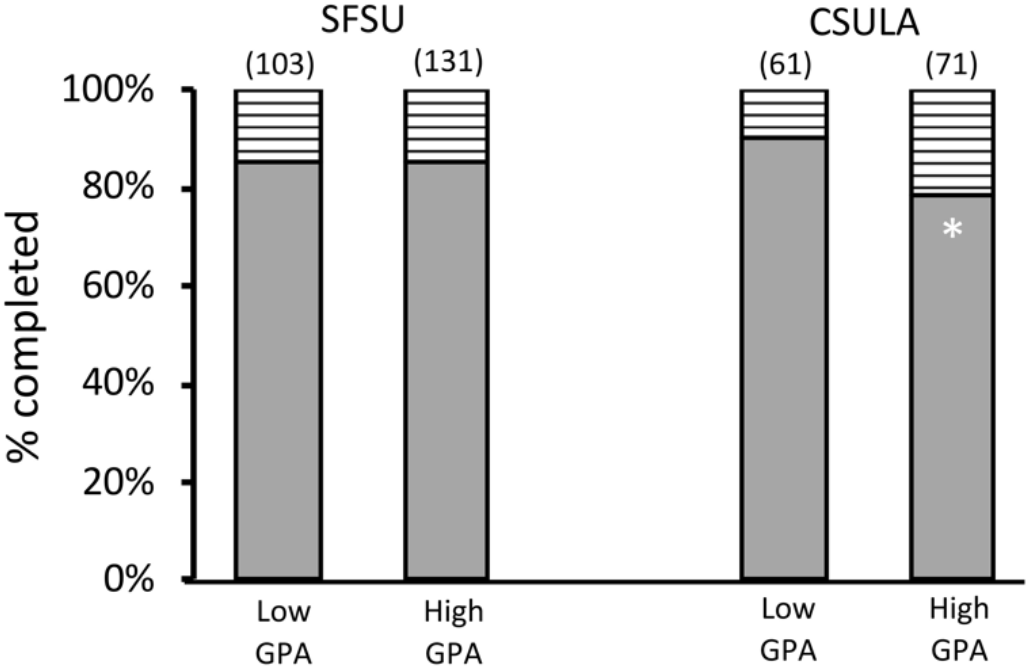
Proportion of SFSU or CSULA students from each GPA group completing a doctoral program. Solid bars represent doctoral programs; striped bars represent incompletes (students in progress are not included here). Total sample sizes are noted above the bars in parentheses. The white asterisk represents a significant difference from the respective Low GPA group (P=0.04).

When pooling all students, so that UG GPA was not considered, the completion rate for all MS students was the same regardless of their undergraduate institution (Table 7). We also considered the likelihood of “in progress” students completing their degrees based on whether their undergraduate degree was obtained from their MS institution or not (Table 8). There was no significant difference in the likelihood of completion for students from SFSU or CSULA (P>0.6). That is, the likelihood of being on track to complete a PhD degree was not dependent on the undergraduate institution.

**Table 7:** Proportion of students from their own or other institutions completing, or still in (“in progress”), a doctoral program.

| Completing PhD<br>% (n) | UG Institution |  | UG Institution |  |
| --- | --- | --- | --- | --- |
|  | SFSU UG | Non-SFSU UG | CSULA UG | Non-CSULA UG |
| Did not complete | 18.1% (15) | 12.8% (19) | 16.3% (7) | 17.6% (16) |
| Completed | 81.9% (68) | 87.2% (132) | 83.7% (36) | 82.4% (75) |
| $\chi^2$ test | 1.3 | | 0.035 | |
| p-value | 0.254 |  | 0.852 |  |
| In progress | 37 | 22 | 8 | 12 |

**Table 8.**
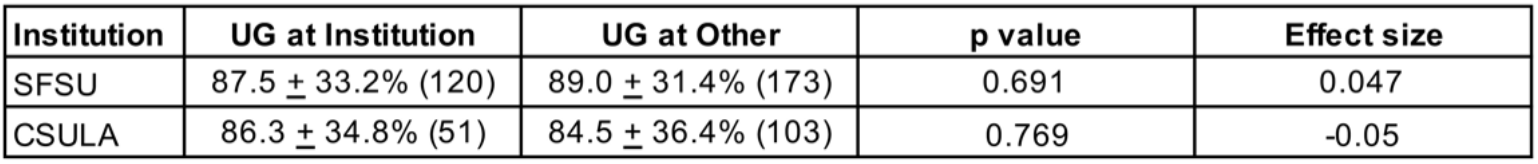
Likelihood of being on track to complete a PhD degree for each undergraduate group.

| Institution | UG at Institution | UG at Other | p value | Effect size |
| --- | --- | --- | --- | --- |
| SFSU | 87.5 ± 33.2% (120) | 89.0 ± 31.4% (173) | 0.691 | 0.047 |
| CSULA | 86.3 ± 34.8% (51) | 84.5 ± 36.4% (103) | 0.769 | -0.05 |

Perhaps most striking in these data were the extremely high completion rates of 80-90% for either cohort, with moderately higher rates for students going to CSULA with low UG GPA levels. This is in sharp contrast to the ∼50% completion rate nationwide (NCSES NSF, 2022).

Finally, of those who already completed a PhD, there was no significant difference in time (measured in years) to complete the degree for SFSU or CSULA MS students (Fig. 4). This was true between the low and high UG GPA groups, as well as when considering which undergraduate institution they came from (Tables 9 & 10). When pooling groups and not considering differences in GPA, UG institution still did not predict a faster time to degree (Table 11).

**Table 9:** Average time for SFSU students from each UG GPA group to complete a doctoral program, sorted by undergraduate institution.

| Time to complete PhD | SFSU |  |  |  |
| --- | --- | --- | --- | --- |
| years (n) | Low GPA (< 3.0) | High GPA ($\geq 3.0$) | p | Effector |
| All Students | 5.92 ± 1.53 (86) | 5.86 ± 1.63 (104) | 0.767 | -0.043 |
| SFSU UG | 5.82 ± 1.77 (25) | 5.62 ± 1.36 (38) | 0.612 | -0.131 |
| non-SFSU UG | 5.97 ± 1.44 (61) | 5.99 ± 1.76 (66) | 0.93 | 0.016 |

**Table 10:** Average time for CSULA students from each GPA group to complete a doctoral, sorted by undergraduate institution.

| Time to complete PhD | CSULA |  |  |  |
| --- | --- | --- | --- | --- |
| years (n) | Low GPA (< 3.0) | High GPA ($\geq 3.0$) | p | Effector |
| All Students | 5.93 ± 1.36 (55) | 5.95 ± 1.53 (56) | 0.945 | 0.013 |
| CSULA UG | 5.67 ± 1.07 (12) | 6.13 ± 1.60 (24) | 0.377 | 0.317 |
| non-CSULA UG | 6.00 ± 1.43 (43) | 5.81 ± 1.49 (32) | 0.583 | -0.129 |

**Table 11:** Average time for all MS students to complete a doctoral program based on whether they went to their own or other UG institutions.

| Institution | Time to degree (n) | p value | Effect size |
| --- | --- | --- | --- |
| SFSU UG | 5.70 ± 1.53 yrs (63) | 0.249 | -0.178 |
| non-SFSU UG | 5.98 ± 1.61 yrs (127) | | |
| CSULA UG | 5.97 ± 1.44 yrs (36) | 0.859 | 0.039 |
| non-CSULA UG | 5.92 ± 1.45 yrs (75) | | |

**Figure 4:**
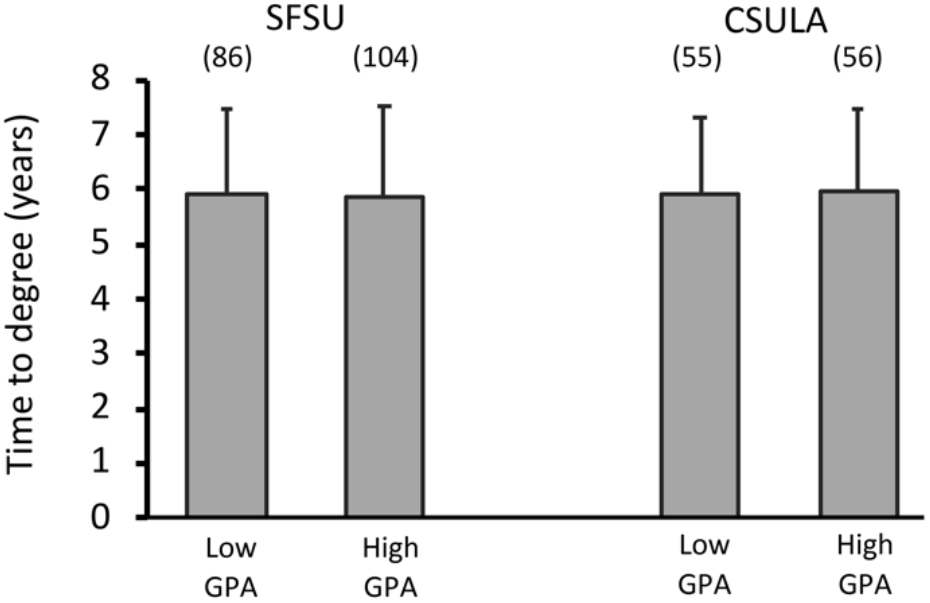
Average time for SFSU or CSULA students from each GPA group to complete a doctoral program. Total sample sizes are noted above the bars in parentheses. Error bars are standard error of the mean.

### 3.4. Inclusive MS training programs can increase student diversity in a Biology Department

Our data suggest that the MS training programs support underrepresented student success in PhD programs independent of UG GPA. Here we provide an example of the positive impact these training grants have had on the MS institution itself.

In the Department of Biology at SFSU, we looked at the overall graduate population in the department, regardless of whether they were funded or not, in each decade (Fig. 5). Groups were identified by race/ethnicity, based on self-reporting (SFSU Institutional Research), where “UR” (underrepresented) represents Native American, African American, Hispanic American, and Pacific Islanders together, as defined by the NIH. “Asian” represents Chinese, Japanese, Korean, and includes South Asian. “Other” represents groups such as white/Caucasian, Middle Eastern, Northern African, and non-disclosed. We found that the total graduate student population almost doubled over the 30 years of data tracking. The increases each decade paralleled the increases in diversity training grants acquired by the SEO office (and therefore the number of funded trainees), as noted by the $$ symbols in Fig. 5. By 2019, there was a 16-fold increase in the UR population. Significantly, this increase did not reduce the “other” population, but it did change the diversity of representation in the department. It should be pointed out that this increased UR population included more than just the funded graduate students, suggesting a positive impact of the funding beyond just the money.

**Figure 5:**
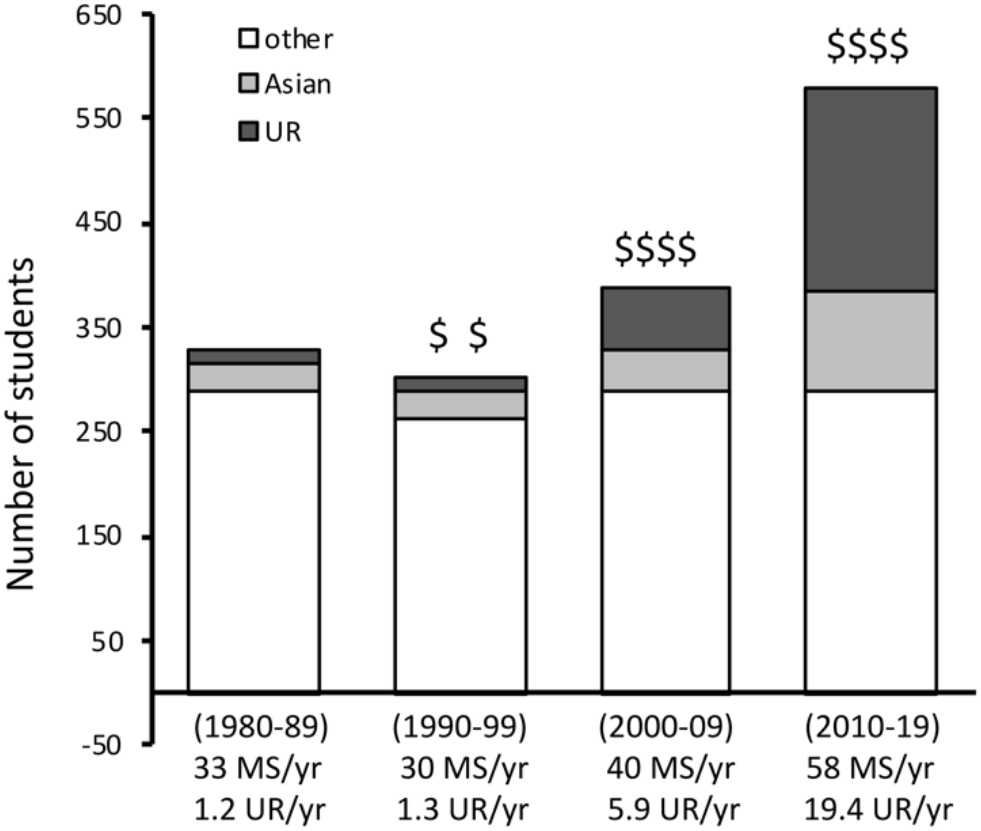
Number of MS students in the Department of Biology at SFSU over 4 decades from 1980-2019, sorted by self-reported race. Diversity training grants are noted by $$ symbols, based on amount of funding. Grants received are listed chronologically as: (i) NIH Bridge to Doctorate, (ii) Graduate Assistance in Areas of National Need (GAANN), (iii) NIH RISE (graduate portion), (iv) NSF LSAMP Bridge to Doctorate (v) NSF Science Technology award for the Center for Cellular Construction, and (vi) Genentech Foundation Scholars Program. “UR” (underrepresented) represents Native American, African American, Hispanic American, and Pacific Islanders together, as defined by the NIH. “Asian” represents Chinese, Japanese, Korean, and includes South Asian. “Other” represents groups such as white/Caucasian, Middle Eastern, Northern African, and non-disclosed.

## 4. Discussion

### 4.1. A high undergraduate GPA does not predict enhanced PhD success

This study looked at 506 students from two California State University (CSU) campuses over a 20-30 year period (1992 to 2019) to assess the importance of undergraduate GPA (UG GPA) in predicting PhD success in the context of participating in funded MS research training programs. The two cohorts of under-represented students who entered PhD programs after participating in the training programs were separated by their UG GPAs (Low UG GPA < 3.0; High UG GPA ≥ 3.0), with a fairly even distribution between the two groups (Table 1). The distribution formed a bell curve from 2.6-3.4 as the majority, where the distribution above a 3.4 GPA was not all that high. In this study, we had three major findings, with only a few caveats: a high undergraduate GPA (≥3.0) did not predict better (1) entry into PhD, (2) completion of a PhD, and (3) time to complete the PhD.

Our first result showed that UG GPA did not predict better success in entry into a PhD program (Fig. 2). This was true whether students came from the same undergraduate institution as their MS institution or another (Tables 2, 3). The rate of student matriculation into these PhD programs ranged from 87-91%. Moreover, undergraduate GPA did not predict the type of PhD institutions attended by students (Tables A1, A2). At SFSU, the top five universities accepting our low UG GPA students were UC Davis, UC Berkeley, University of Washington, UCSF, and Harvard University. The top five universities accepting our high UG GPA group were UC Davis, UCSF, UC Berkeley, UCLA, and Harvard University, all R1 institutions, with strong reputations. The top five PhD institutions accepting CSULA students from the low UG GPA group were UCLA, UC San Diego, UC Riverside, UC Davis, and UC Irvine/University of Illinois, Urbana-Champaign (equal numbers). The top five universities accepting CSULA students from the high UG GPA group were UCLA, University of Southern California, UC Riverside, UC Irvine, and Claremont Graduate University.

Our second result indicated that a high UG GPA did not predict better PhD completion rates (Fig. 3). In fact, CSULA MS students who came from another undergraduate institution had a significantly higher completion rate, at over 90%, when they had an undergraduate GPA *less* than 3.0 (Table 6). This is even stronger evidence that high UG GPA does not accurately predict success, and it would be interesting to examine whether other factors, such as research experience, vary between these groups. Perhaps a more important result of this study that is worth pointing out is the high overall completion rate, at over 80%. This is far above the national average of ∼50% in the biomedical sciences (NCSES NSF, 2022). We assessed the “In Progress” students, and likewise found a high likelihood of completing their PhDs at a rate of over 80% (Table 8).

Finally, we found that a high UG GPA did not predict a faster time to complete the PhD (Fig. 4). The results are incredibly similar between the low and high UG GPA groups, typically at under 6 years for both institutions (Tables 9-11). This aligns well with the national average for the biomedical sciences at 5.7 years (NSF, 2021).

When we did not consider undergraduate GPA, success was also equal, between students staying at their undergraduate institution for their MS compared to those coming from another undergraduate institution. There was no significant difference in the matriculation, completion or time to degree for these students (Tables 4, 7 and 11).

### 4.2. MS diversity training programs housed in “centers” as vital interventions for the PhD pipeline

IMSD programs across the country often tout the “top” universities from which their PhD students come as a metric of the quality of their students. We suggest here that the institution does not matter as much as the quality of support it gives its students and its ability to provide an intervention for under-represented students struggling with personal issues such as stereotype threat, imposter syndrome, financial hardships and lack of counseling in academia. This argument is bolstered by the fact that our PhD completion rates averaged over 80%, far above the national average of ∼50% in the biomedical sciences (NCSES NSF, 2022).

The California State University (CSU) system needs to be considered as a major pathway for diversifying the PhD. It is the largest public university system in the United States, comprised of 23 campuses and 7 off-campus centers, enrolling almost 500,000 students and employing over 50,000 faculty and staff members (Wikipedia. “California State University”, 2024). CSU is noted for its diversity, educating some of the nation’s most racially, economically, and academically varied student populations (Grove, 2023), and it houses many of the training grants offered by the NIH and NSF along with private corporations. Moreover, grade inflation has been slower in the CSU system than at more selective colleges (Carter, 2016), an observation consistent with earlier studies of similar comprehensive state systems. CSU admissions are only modestly selective and most of its urban campuses serve largely commuter populations of students (Rojstaczer, 2010; Rojstaczer 2012). Both SFSU and CSULA are minority serving institutions. Within SFSU, the diversity of under-represented students is approximately 48% (including two or more races), with 31% first-generation and 70% receiving financial aid (SFSU Institutional Research). At CSULA, under-represented students comprise 80% of the enrollment. Eighty-five percent are in the first generation in their families that will earn baccalaureate degrees, 83% receive financial aid and 63% are eligible for Pell Grants (CSULA Institutional Effectiveness).

We suggest that participation in research training programs such as those funded by NIH or NSF, can make UG GPA as a predictor to PhD success irrelevant. The MS training grants, which include a strong independent research experience coupled with directed professional development and multifaceted mentorship, are strong mechanisms for the success of diversity initiatives. We further suggest that we need to evaluate the effectiveness of “offices” or “centers” to house multiple training grants as a mechanism to maximize the limited administrative funding provided by training grants, where programs can be tiered, and near-peer mentoring can be implemented from Community Colleges through to MS programs.

Finally, we provide an example of the impact of these MS training grants on the MS programs themselves, providing a strong positive-feedback loop (Fig. 5). The total number of graduate students in the Department of Biology at SFSU increased proportionately to the number of funded MS training grants acquired by the SEO office. After 30 years, the number of students nearly doubled, but UR students did not take the spots of non-UR students. Rather, they enhanced the diversity of the department. That is, the total number of non-UR students did not change, but within the last decade there was a 16-fold increase in UR students. Interestingly, this increase included both funded and non-funded students, highlighting the broader impacts of these grants in providing the improved environment for diverse student success. We suggest this was due to a combination of increased UR presence in the department to attract more UR students to the department, as well as the synergistic activities between the SEO office training grants and the Department of Biology MS curriculum (e.g. The introductory Biology course for all incoming MS Biology students was in part modeled after the training curriculum from programs such as the NIH Bridge to Doctorate training grant).

## 5. Conclusion

The holistic approach we used to admit students into our funded MS research-training programs, and the developmental opportunities they offer them have been successful in preparing students for success in PhD studies. The MS research training programs include financial support (stipends, tuition, conference travel and research supplies) and academic support (e.g. professional development workshops/courses), and a mentoring ecosystem for students that includes program directors, program staff, class instructors, research advisors, and near-peer mentors, along with community-building activities). Review of candidates included letters of recommendation, personal statements, prior research experience, with less emphasis on undergraduate transcripts. All these inputs became the context for interviews with the candidates, among whom was a sizeable population that was highly recommended and had considerable research experience, but, surprisingly, lower U GPAs (2.0-2.9). We suggest that the lower UG GPA did not reflect a lack of talent, but more likely the results of (i) working too much while attending college, (ii) little academic support for first-generation students, and/or (iii) high levels of stereotype threat and imposter syndrome (Weinstein, 2002; Moss-Racusin *et al*., 2012; Leslie *et al*., 2015; Estrada *et al*., 2016).

Top research doctoral institutions fundamentally ask applicants to their PhD programs: “Can you learn to do research really well? Are you inquisitive, motivated, and prepared to do this?”. We maintain that undergraduate GPA is a poor proxy for talent and motivation for a career in research. It is not that high grades cannot, in conjunction with letters of recommendation, statements of purpose, prior research experience, and interviews, identify good candidates for PhD programs. Clearly, they can. The more significant question is: which highly talented candidates are missed by using GPA as an early filter, rather than as one among several factors in a holistic evaluation of applicants for admission?

