## Supplement 1 for "Undergraduate GPA does not predict success in PhD programs for cohorts of MS students at two minority-serving institutions"

**Table S1. Distribution of students by GPA at SFSU or CSULA.**

| GPA Range | SFSU | CSULA |
|---|---|---|
| | % (n) | % (n) |
| 2.00-2.19 | 3 (2) | 1 (2) |
| 2.20-2.39 | 6 (4) | 3 (5) |
| 2.40-2.59 | 22 (14) | 5 (8) |
| 2.60-2.79 | 43 (28) | 14 (22) |
| 2.80-2.99 | 69 (45) | 19 (29) |
| 3.00-3.19 | 60 (39) | 21 (32) |
| 3.20-3.39 | 43 (28) | 13 (20) |
| 3.40-3.59 | 30 (19) | 8 (12) |
| 3.60-3.79 | 20 (13) | 9 (14) |
| 3.80-4.00 | 10 (6) | 7 (11) |
