## Supplement 2 for "Undergraduate GPA does not predict success in PhD programs for cohorts of MS students at two minority-serving institutions"

**Table S2: Institutions attended by 5 or more SFSU MS students in PhD programs over 30 years, sorted by low and high undergraduate GPA levels. Completion rates are lower than entry rates, since some students are still in progress and some (<7%) withdrew.**

| PhD Institution | SFSU Low GPA | | SFSU High GPA | | SFSU Total | |
|---|---|---|---|---|---|---|
| | Enter | Complete | Enter | Complete | Enter | Complete |
| University of California Davis | 26 | 19 | 22 | 15 | 48 | 34 |
| University of California San Francisco | 8 | 4 | 14 | 10 | 22 | 14 |
| University of California Berkeley | 9 | 6 | 10 | 9 | 19 | 15 |
| Harvard University | 8 | 4 | 7 | 5 | 15 | 9 |
| University of California Los Angeles | 3 | 2 | 10 | 7 | 13 | 9 |
| Stanford University | 6 | 2 | 6 | 5 | 12 | 7 |
| University of Washington | 9 | 5 | 3 | 1 | 12 | 6 |
| University of California Santa Cruz | 6 | 4 | 4 | 2 | 10 | 6 |
| University of Michigan | 4 | 2 | 5 | 3 | 9 | 5 |
| University of California San Diego | 2 | 1 | 6 | 4 | 8 | 5 |
| University of Texas, South Western | 5 | 3 | 2 | 2 | 7 | 5 |
| University of California Irvine | 3 | 1 | 4 | 2 | 7 | 3 |
| New York University | 4 | 4 | 2 | 1 | 6 | 5 |
| University of Wisconsin-Madison | 1 | 1 | 4 | 4 | 5 | 5 |
| **TOTAL** | **94** | **58** | **99** | **70** | **193** | **128** |

* OTHER institutions (in order of number of acceptances) include: Albert Einstein School of Medicine, Utah State U., U. Arizona, Washington U., Emory U., U. Florida, U. Southern California, Johns Hopkins U., Northwestern U., U. Maryland, U. Montana, U. Tennessee, U.T. San Antonio, Washington State U., Arizona State U., Baylor Medical College, Boston U., Brown U., Texas A&M, U. South Florida, U. Utah, U.C. Merced, U.N.C. Chapel Hill, California Institute of Technology, Cambridge U., College of William & Mary, Columbia U., Cornell U., Dartmouth U., Drexel U., Eastern Virginia Medical School, Kobe U., Loma Linda U., Massachusetts College of Pharmacology, Memorial Sloan Kettering, Pennsylvania State U., Portland State U., Ross U., Seton Hall U., State U. of New York Stony Brook, Temple U., Tulane U., Université Cote d'Azur, U. Alabama, U.C. Riverside, U.C. Santa Barbara, U. Chicago, U. Georgia, U. Hawaii, Manoa, U. Illinois, Urbana-Champaign, U. Iowa, U. Melbourne, U. Missouri, U.T. Health Center Houston, U.T. Austin, U. Pacific, U. Virginia, US Air Force Institute of Technology, Victoria University of Wellington, City University of New York, Florida State U., George Mason U., Georgia Institute of Technology, HHS Center for Disease Control, U. Buffalo, U. Colorado Boulder, U. Nevada Reno, U. North Texas, U. Oregon, Virginia Tech U., Yale U.
