## Supplement 3 for "Undergraduate GPA does not predict success in PhD programs for cohorts of MS students at two minority-serving institutions"

**Table S3: : Institutions attended by 5 or more CSULA MS students in PhD programs over 20 years, sorted by low and high undergraduate GPA levels. Completion rates are lower than entry rates, since some students are still in progress and some (<20%) withdrew.**

| PhD Institution | CSULA Low GPA | | CSULA High GPA | | CSULA Total | |
|---|---|---|---|---|---|---|
| | Enter | Complete | Enter | Complete | Enter | Complete |
| University of California Los Angeles | 8 | 4 | 14 | 10 | 22 | 14 |
| University of Southern California | 3 | 3 | 12 | 5 | 15 | 8 |
| University of California Riverside | 5 | 4 | 5 | 3 | 10 | 7 |
| University of California Irvine | 4 | 4 | 5 | 4 | 9 | 8 |
| University of Illinois, Urbana-Champaign | 4 | 4 | 4 | 2 | 8 | 6 |
| University of California San Diego | 6 | 6 | 1 | 1 | 7 | 7 |
| University of California Santa Cruz | 2 | 1 | 4 | 1 | 6 | 2 |
| University of Washington, Seattle | 3 | 3 | 3 | 2 | 6 | 5 |
| Claremont Graduate University | 0 | 0 | 5 | 2 | 5 | 2 |
| University of California Davis | 5 | 2 | 0 | 0 | 5 | 2 |
| **TOTAL** | **40** | **31** | **53** | **30** | **93** | **61** |

* OTHER institutions (in order of number of acceptances) include: City of Hope National Medical Center, University of Michigan, Ann Arbor, New Mexico State U., Stanford U., U. Arizona, U.C. Berkeley, U.C. Merced, U. Colorado Boulder, U.N.C. Chapel Hill, U. Pittsburgh, Alabama State University Tuscaloosa, Albert Einstein School of Medicine, Arizona State U., Azusa Pacific U., Baylor U., California Institute of Technology, Columbia U., Cornell U., Georgia Institute of Technology, Georgia State U., Harvard U., Indiana University School of Medicine, James Cook University (Australia), Louisiana State U., Massachusetts Institute of Technology, Utah State U., U. Arizona, Washington U., Emory U., U. Florida, U. Southern California, Johns Hopkins U., Max Planck Institute for Astronomy (Germany), Michigan State U., Mississippi State U., New York U., Northeastern U., Sackler Institute for Comparative Genomics, American Museum of Natural History, Scripps Research Institute, La Jolla, Temple U., Texas A&M, U. Alabama, Birmingham, U. Cincinnati, U. Colorado, Denver, U. Florida, U. Kentucky, U. Massachusetts Amherst, U.T. Austin, U. Wisconsin, Madison, Vanderbilt U., Western Michigan U., Yale U.
